# Neuronal correlates of attentional selectivity and intensity in visual area V4 are invariant of motivational context

**DOI:** 10.1101/2021.07.30.454515

**Authors:** Supriya Ghosh, John H.R. Maunsell

## Abstract

Flexibly switching attentional strategies is crucial for adaptive behavior in changing environments. Depending on the context, task demand employs different degrees of the two fundamental components of attention– attentional selectivity (preferentially attending to one location in visual space) and effort (the total non-selective intensity of attention). Neuronal responses in the visual cortex that show modulation with changes in either selective attention or effort are reported to partially represent motivational aspect of the task context. The relative contributions and interactions of these two components of attention to modulate neuronal signals and their sensitivity to distinct motivational drives are poorly understood. To address this question, we independently controlled monkeys’ spatially selective attention and non-selective attentional intensity in the same experimental session during a novel visual orientation change detection task. Attention was controlled either by adjusting the relative difficulty of the orientation changes at the two locations or by the reward associated with stimuli at two locations while simultaneously recording spikes from populations of neurons in area V4. We found that V4 neurons are robustly modulated by either selective attention or attentional intensity. Notably, as attentional selectivity for a neuron’s receptive field location decreased, its responses became weaker, despite an increase in the animal’s overall attentional intensity. This strong interaction between attentional selectivity and intensity could be identified in single trial spike trains. A simple divisive normalization of spatially distributed attention performances can explain the interaction between attention components well at the single neuron level. The effects of attentional selectivity and attentional intensity on neuronal responses were the same regardless of whether the changes were motivated by reward or task difficulty. These results provide a detailed cellular-level mechanism of how fundamental components of attention integrate and affect sensory processing in varying motivational and stimulus contexts.

## INTRODUCTION

Attention plays an essential role in motivating human behavior and cognition by selectively enhancing the processing of relevant sensory information. To engage and perform in cognitively demanding tasks, goal directed attention is often driven by external incentives. Many cortical and subcortical brain areas, including V4, change their activity when attention shifts^1-4^. They are also sensitive to the size of the reward that motivates those shifts^5-11^. Although reward expectation and attention have been described as conceptually distinct cognitive constructs (for review^12^), it remains challenging to distinguish these factors owing to their covariance and the high similarity of their effects on neuronal responses. Attentional levels can also be elevated owing to internal desire to complete a task without any apparent changes in external incentives, such as increased cognitive demand as a result of increased task difficulty^13^. For example, professional athletes or musicians address demanding situations with increased effort so as to maintain a given performance level. The contributions of different sources of motivation to regulation of sensory processing in cortex and overall perceptual behavior remain elusive.

In order to adapt to varying environmental and stimulus contexts, subjects shift their attention between spatially localized targets or selective stimulus features to spatially global targets or nonselective features. Many studies have characterized the neuronal modulations associated with selective attention by assaying how performance improves for attended spatial locations or stimulus features relative to distant locations or unrelated features. When a monkey’s attention is selectively directed towards the location of a neuron’s receptive field (RF), improvement in perceptual performance in that region is typically accompanied by increased spike rates^4,14^, reduced individual response variance and pairwise spike count correlations^1,15^. Although experimental studies most often treat attention as all-or-none, it has another fundamental aspect, intensity^16^–how strongly attention is focused independent of selectivity. Attentional intensity can be considered as an objective measure of perceived effort or cognitive engagement in a goal directed attention demanding task^17^. Attention related modulations of neurons in area V4 in primate visual cortex have been examined using a variety of visual detection tasks^2,3,18,19^. Some of these studies show that V4 neuronal activity is enhanced as a result of higher cognitive engagement or attentional effort in response to increased task demand^13,20^. It remains relatively unknown how selective attention and attentional intensity integrate in the brain to improve sensory perception and performance, and how motivational contexts influence these processes.

To address these questions, we trained monkeys to do an attention demanding visual task that allowed us to independently control the monkey’s attentional selectivity and intensity in two different motivation contexts. We varied either task difficulty while reward size was kept fixed or varied reward size for a fixed task difficulty. Using simultaneous electrophysiological recordings from populations of V4 neurons and computational models, we found that attentional selectivity and intensity independently modulate neuronal spiking. Single trial spike trains encode multiplexed signals of attentional selectivity and intensity with comparable strengths in a way that is independent of the how the animal was motivated to allocate its attention. Further, the effects of attentional selectivity and intensity interact to determine the resultant influence of attention on spiking. A spatially tuned normalization model of attention can account for this interaction. Thus, we provided a detailed account of how fundamental components of attention interact at the level of V4 spikes. By extending the spectrum of attention-related cognitive representations in V4, the result provide help clarify how individual neurons contribute to higher-order cognition.

## RESULTS

### Independent control of attentional selectivity and intensity by varying task difficulty

We trained two rhesus monkeys to distribute their visual spatial attention between stimuli in the left and right hemifields while doing an orientation change detection task (**Figure 1a**). The animal held its gaze on a central fixation spot throughout each trial. After a randomly varying period of fixation, two Gabor sample stimuli appeared for 200 ms. This was followed by a delay of 200-300 ms, after which a single Gabor test stimulus appeared at one of the two sample locations (selected pseudo-randomly). If the orientation of the test stimulus differed from the orientation of the sample stimulus that had appeared in that location (a target), the monkey had to rapidly make a saccade to the stimulus to earn a juice reward. On a random 50% the trials, the orientation of test stimulus was unchanged (a non-target) and the monkey was required to maintain fixation. In that case, a second test stimulus that always had a different orientation was presented after a short delay and monkey needed to saccade to this target stimulus to earn a reward.

**Figure 1.**
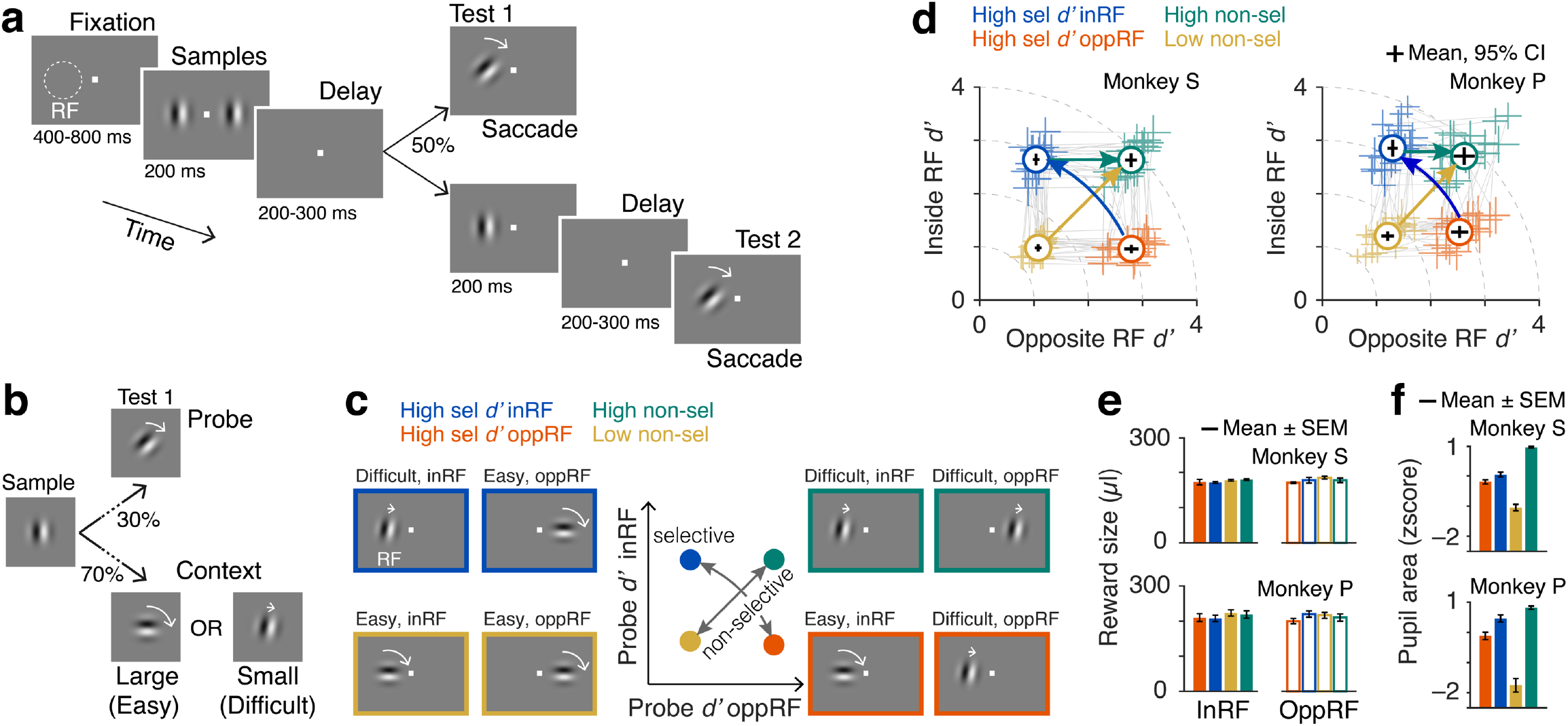
Independent control of attentional selectivity and intensity using differential task demands. **a** Visual spatial attention task. Monkeys were required to fixate, attend to sample stimuli (Gabors) presented in both hemifields (inside and opposite side of recorded neurons’ receptive field (RF)) and report an orientation change that occurred in one of the two test intervals by making a saccade to the stimulus location. **b** Task demands. Orientation changes on ∼30% of the non-matched trials are intermediate (probe) and on the rest ∼70% of the non-matched trials are either large (easy context) or small (difficult context). **c** Centre, distribution of task difficulties across two locations in opposite hemifields for four task conditions: high selective attention either inside the RF (blue) or opposite RF location (orange) when attentional intensity remains fixed, and low or high non-selective attentional intensity (yellow and green). Left and Right, Four attention conditions consisted of different combinations of these task contexts (easy and difficult) at the two stimuli locations. **d** Attention operating characteristic (AOC) curve, indicating behavioral sensitivity (d’) on individual sessions and their average (circles) for test stimuli inside and opposite side RF during the four attention conditions, (sessions: 20 monkey S; 22 monkey P). Grey lines connect attention conditions within a session. Dotted lines, iso-intensity lines. Error bars, 95% confidence intervals. **e** Session averaged reward sizes across different attention conditions for individual monkeys. **f** Session averaged pupil area (z-scored) during the sample stimuli period. Error bars, ± SEM.

To control the animal’s attention, in each block of trials we set the orientation change of the first test stimulus at each location to be either easy to detect (∼80°) or difficult to detect (∼18°) (**Figure 1b**). In each block the size of the orientation change at the two locations was set independently, providing four possible combinations (**Figure 1c**). We measured the behavioral consequences of different combinations of difficulty by presenting an orientation change of intermediate difficulty (30°, probe) on a randomly selected fraction of all trials (∼30%). These probe trials allowed us to directly compare behavioral sensitivity (*d’*) at both location across all four block types. **Figure 1d** plots the average *d’*s for the left and right stimulus locations on probe trials for the two monkeys separately, with different colors representing the four different combinations of difficulty. Crosses mark mean *d’*s from individual sessions and gray lines join the four means from individual sessions. The changes in behavioral performance document that the animals responded to task difficulty by adapting their allocation of attention. Behavioral *d’* for the probe orientation change on each side was substantially higher when most orientation changes were difficult to detect, and lower when most changes were easy to detect, with approximately symmetrical *d’*s at both locations during most individual sessions.

Spatial selectivity of attention was quantified by selectivity index that measured the relative behavioral *d’* at the RF location compared to the opposite location (**Methods**). Attentional intensity was measured by overall absolute behavioral *d’*s in the two locations (**Methods**). The ∼4-fold difference in orientation change (median easy change 80°, IQR 80°-90°; median difficult change 18°, IQR 16°-18°) strongly motivated animals to adjust their behavioral *d’*, whether the inter-block changes on the two sides were in opposite directions (blue arrows, **Figure 1d**) or in the same direction (gold arrows, **Figure 1d**). In both cases behavioral *d’* changed by ∼2-fold (*selectivity indices* in the opposite direction: monkey S, for low RF *d’* mean –0.58 SEM 0.01; for high RF *d’* 0.52 mean 0.01 SEM, p < 10^−12^; monkey P, for low RF *d’* mean –0.41 SEM 0.02; for high RF *d’* mean 0.46 SEM 0.01; p < 10^−13^; *attentional intensity* in the same direction: monkey S, for non-selective low *d’* mean 1.46 SEM 0.04; for non-selective high *d’* mean 3.84 SEM 0.08; p < 10^−9^; monkey P, for non-selective low *d’* mean 1.71 SEM 0.07; for non-selective high *d’* mean 3.78 SEM 0.11; p < 10^−13^; **Supplementary Table T1**). These changes in allocation of attention were driven by changes in task difficulty alone. Although reward sizes for correct responses were varied somewhat from trial to trial (see below), the average was kept the same on both sides across all task difficulty configurations (**Figure 1e**).

Non-luminance mediated task-evoked increases of pupil size are commonly considered a proxy for arousal or attentional engagement and are sensitive to task demands across species^21-23^. Consistent with this, pupil area during the sample stimuli increased progressively with the increase in attention intensity (F_(3, 76)_ = 62.71, p < 10^−19^ for monkey S; F_(3, 84)_ = 88.19, p < 10^−25^ for monkey P, ANOVA; **Figure 1f**). Pupil area was greatest when discriminations were difficult on both sides, and smallest when they were easy on both sides.

### Relative neuronal modulation of V4 with attentional selectivity and non-selective intensity

We recorded from 1194 single units and small multi-unit clusters (single unit, 385; multiunit, 809) during 42 recording sessions from the two monkeys (monkey S, 20 sessions, 714 units; monkey P, 22 sessions, 480 units) using 96 channel multielectrode arrays chronically implanted in V4 in the superficial prelunate gyrus. Neurons typically responded more strongly to the sample stimuli during the trial blocks when the monkey’s behavioral *d’* at the RF location was high. This increased spiking response was seen whether the behavioral *d’* differences involved different selectivity for the RF versus other location (red versus blue) or a change in attentional intensity with no change in selectivity (gold versus green). Importantly, the spike responses did not depend exclusively on *d’* in the RF location. Neuronal responses during high non-selective behavioral *d’* (green, **Figure 2a-b**) were reduced compared to responses with identical RF *d’* and low d’ at the distant location (blue, **Figure 2a-b**).

**Figure 2.**
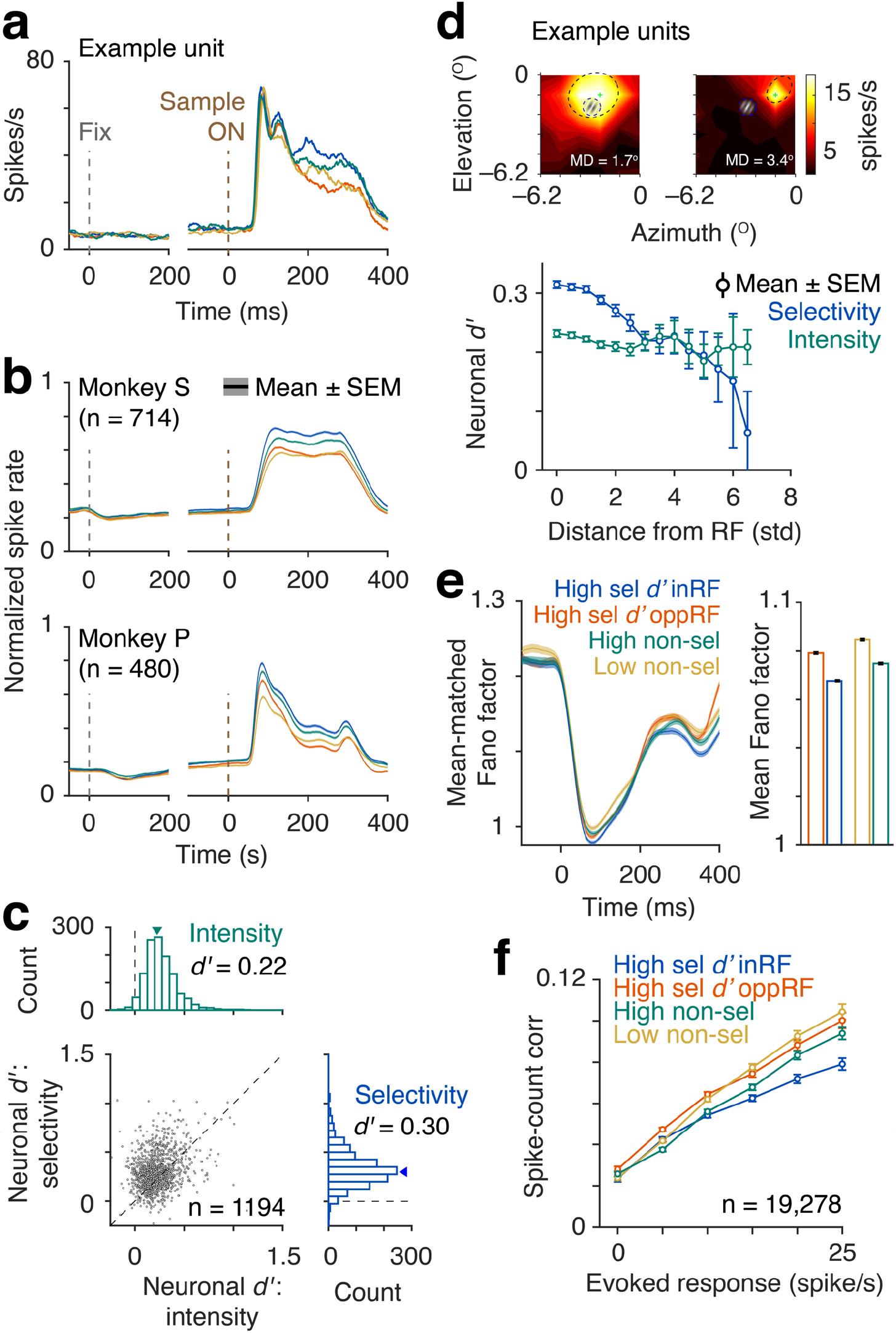
Neuronal modulation with changes in attentional selectivity and intensity. **a** Peri-stimulus time histograms (PSTH) of spike rates of correct trials in different attention conditions for an example neuron in V4. Single trial spike counts were binned at 2 ms, smoothed with σ = 15 ms half-Gaussian and then aligned at the onset of sample stimulus. **b** Population PSTHs for monkey S (top) and monkey P (bottom). For population average, spike rates of each neuron were normalized to its peak response within 60 - 260 ms from sample stimulus onset (monkey S, n = 714; monkey P, n = 480). **c** Distribution of neuronal d’ for attentional selectivity and intensity of all units from both monkey S and P (n = 1194). **d** Top, Receptive field (RF) locations of two example units relative to the Gabor stimulus. MD, Mahalanobis distance measures the standardized distance between neurons’ RF and Gabor stimulus in units of standard deviation. Bottom, Distribution of neuronal d’ as a function of RF-Gabor distance. **e** Left, Mean-matched Fano factor. Right, Mean-matched Fano factor averaged over 60-260 ms from the sample onset. Error bars, ±SEM. **f** Pairwise correlations between spike-counts of simultaneously recorded neurons over 60 to 260 ms from sample onset (n = 19,278 pairs, all units) and binned according to their evoked responses (geometric mean of baseline subtracted spike counts). Error bars, ±SEM.

To quantify neuronal modulation by attentional selectivity and intensity, we computed a neuronal *d’* as the difference of z-scored firing rates (60-260 ms from sample onset) between high and low attention states. The mean firing rate modulation was significantly greater for attention selectivity compared to non-selective intensity (neuronal *d’* for selective attention, mean±SEM = 0.30±0.01; for non-selective attention, mean ± SEM = 0.22 ± 0.05, p < 10^−31^, *n* = 1194, t-test; **Figure 2c**). Single neurons and multiunit clusters separately showed similar spike modulation by attentional selectivity and non-selective intensity (single units, p < 10^−7^; multiunits, p < 10^−26^). These attentional effects were also significant for the monkeys individually (monkey S, p < 10^−27^; monkey P, p < 10^−8^).

Changes in attentional intensity with this task design do not rule out all forms of spatial selection because the animals might have attended to locations other than the two stimulus locations tested. If so, V4 neurons could have been modulated by the spatially selective shifting of attention from those other locations to the two stimulus locations. We examined the broader spatial distribution of attention by measuring the correlation between firing rate modulation and the proximity of a V4 neuron’s RF and the attended Gabor stimulus (**Figure 2d**) using Mahalanobis distance to measure proximity. As expected, neuronal *d’* dropped substantially with increasing RF distance from the stimulus center when animals shifted their spatially selective attention (Spearman, *ρ* = –0.18, p < 10^−8^; **Figure 2d**). In contrast, *d’* for the same neurons varied little with RF distance when animals were encouraged to adjust their attentional intensity (Spearman, *ρ* =– 0.05, p = 0.11; **Figure 2d**), supporting the absence of spatially selective attention in this manipulation. Correlation between the RF distance and neuronal *d’* for attentional selectivity was significantly higher compared to attentional intensity (p = 0.002, z-test).

Because we recorded from the same fixed multielectrode arrays over many sessions, it is possible that some units were sampled in more than one session. We investigated the effect of potential resampling by analyzing a subsample that included only one unit from each electrode across all recording sessions (n = 85 for monkey S, n = 80 for monkey P). For this conservative set of unequivocally unique units, both the high selective and non-selective intensity increased spike rates and the modulation was stronger for selective than non-selective attention (mean ± SEM neuronal *d’* for monkey S, selective attention, 0.25 ± 0.01, non-selective attention, 0.19 ± 0.01, p < 10^−3^; for monkey P, selective attention, 0.31 ± 0.02, non-selective attention, 0.25 ± 0.02, p = 0.02, t-test; **Supplementary Figure S2**) by amounts that were indistinguishable from the whole population. Thus, the results cannot be attributed to multiple sampling that might have occurred from units with uncharacteristic properties.

In addition to spike rate modulation, we tested the relative effects of attentional selectivity and non-selective intensity on signal-to noise of individual V4 units by measuring mean-matched Fano factor (the ratio of the variance of the spike counts to the mean). Fano factors during the sample stimulus period were significantly reduced by increased attention selectivity (F_(1, 140996)_ = 236.82, p < 10^−10^, ANOVA) as well as intensity (F_(1, 140996)_ = 705.29, p < 10^−10^, ANOVA; **Figure 2e**). Further, a significant interaction was detected between the selectivity and intensity on the Fano factor (F_(1, 140996)_ = 4.63, p = 0.03, ANOVA). Similar to the signal-to-noise, pairwise spike count correlations of simultaneously recorded units were also reduced with higher attention selectivity and intensity (selectivity, F_(1, 124917)_ = 16.04, p < 10^−3^; intensity, F_(1, 124917)_ = 167.04, p < 10^−10^; interaction, F_(1, 124917)_ = 7.12, p = 0.008, ANOVA; **Figure 2f**). These results suggest that although the neuronal modulation by selective attention is stronger than the modulation by non-selective intensity, they share many similarities and both contribute appreciably to attention-related modulations.

### Independent control of attentional selectivity and intensity using differential reward size as the external motivator

Subjects are motivated in many different ways to allocate their attention. So far in our task, changes in task difficulty motivated monkeys to spatially redistribute their attention in order to match task demands. We next tested whether the encoding of attention components in V4 neurons depends on how animals are motivated to attend. For this, we instructed the same monkeys to shift their spatial attention by varying reward sizes between the two locations (**Figure 3a-b**). The size of the orientation change was kept constant and challenging on both sides throughout the session. Consequently, no probe orientation changes were needed or presented. The reward manipulation sessions were conducted on different days that were interleaved with the task difficulty sessions described in previous sections. The trial distributions and the probability of an orientation change in the two locations in opposite hemifields were same. Thus, the allocation of spatial attention across the hemifields were primarily motivated by the reward distributions. **Figure 3c** plots behavioral *d’* on the first test stimuli on the left and right sides for all four reward schedules. Behavioral *d’*s on both locations were symmetrical during most individual sessions.

**Figure 3.**
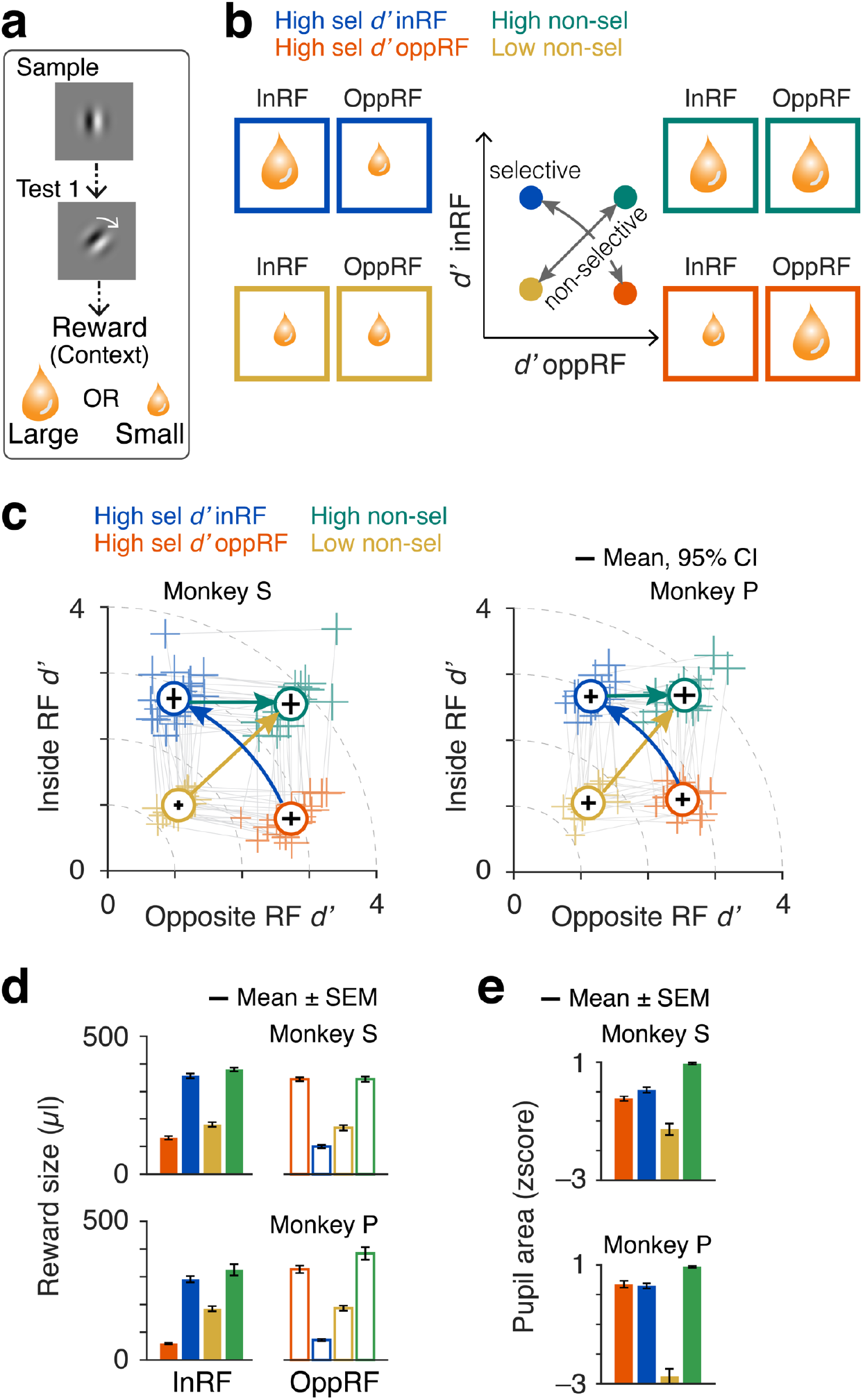
Independent control of attentional selectivity and intensity using differential reward size. **a** Task demands. Orientation changes on ∼30% of the non-matched trials are intermediate (probe) and on the rest ∼70% of the non-matched trials are either large (easy context) or small (difficult context). **b** Centre, distribution of task difficulties across two locations in opposite hemifields for four task conditions: high selective attention either inside the RF (blue) or opposite RF location (orange) when attentional intensity remains fixed, and low or high non-selective attentional intensity (yellow and green). Left and Right, Four attention conditions consisted of different combinations of these task contexts (easy and difficult) at the two stimuli locations. **c** Distribution of reward sizes across two stimuli locations (inRF and oppRF) for four attention conditions. Attention conditions consist of different combinations of reward sizes at the two stimuli locations. **d** Session averaged reward sizes across the four conditions for individual monkeys. **c** Attention operating characteristic (AOC) curve, indicating behavioral sensitivity (d’, circles) on individual sessions and their average (solid markers) for test stimuli inside and opposite side RF during (sessions: 20 monkey S; 16 monkey P). Dotted colored lines indicate average d’ in each hemifield. Lines connect two reward conditions within a session. Error bars, 95% confidence intervals. **e** Session averaged pupil area (z-scored) sample stimuli periods. Error bars, ± SEM.

The ∼2.5-fold increase in reward size (median small 136 μl, IQR 87-177 μl; median large 340 μl, IQR 305-373 μl; **Figure 3d**) strongly motivated animals to adjust their behavioral *d’*, whether the direction of change on the two sides was opposite (blue arrows, **Figure 3c**) or the same (gold arrows, **Figure 3c**). In both cases behavioral *d’* changed by ∼2-fold (opposite direction: selectivity indices in RF, monkey S, for low *d’* mean –0.64 SEM 0.02, for high *d’* 0.54 mean 0.02 SEM, p < 10^−11^; monkey P, low *d’* mean –0.47 SEM 0.02; high *d’* mean 0.48 SEM 0.02; p < 10^− 11^; same direction: intensity indices, monkey S, for low non-selective intensity mean 1.45 SEM 0.05; for high non-selective intensity mean 3.72 SEM 0.1; p < 10^−9^; monkey P, for low non-selective intensity mean 1.53 SEM 0.07; for high non-selective intensity mean 3.70 SEM 0.09; p < 10^−10^). These changes in attention allocation were produced by changes in reward size alone.

As with changes in task difficulty, high non-selective attention intensity mediated by reward size was also associated with increased pupil area during the sample stimuli (F_(3, 76)_ = 54.12, p < 10^−18^ for monkey S; F_(3, 60)_ = 160.47, p < 10^−27^ for monkey P; ANOVA; **Figure 3e**).

### Neuronal modulation of attention selectivity and intensity by differential reward size was indistinguishable from task difficulty mediated modulation

We recorded from total 1331 single units and small multi-unit clusters (single unit, 419; multiunit, 912) in V4 during 36 reward manipulation recording sessions from the two monkeys (monkey S, 20 sessions, 850 units; monkey P, 16 sessions, 481 units). Spike response modulation was similar to the effects observed when attention was controlled using task difficulty (**Figure 4a-c**). Population PSTHs for correctly completed trials increased with high selective attention inside the neuron’s RF (orange and blue traces, **Figure 4a-b**). Spiking activity also increased for higher non-selective attention intensity but was relatively smaller compared to the modulation due to increased selective attention (yellow and green traces, **Figure 4a-b**). Neuronal *d’* for attention selectivity was higher than non-selective intensity (mean ± SEM for selectivity, 0.37 ± 0.006; intensity, 0.24 ± 0.004; p < 10^−72^, t-test) and they were significantly correlated (*ρ* = 0.25, p < 10^−19^, Spearman; **Figure 4c**). Compared to the non-selective intensity, neuronal *d’* for attention selectivity dropped more strongly compared to non-selective intensity with the RF-sample stimulus (selective attention, *ρ* = –0.21 (p < 10^−13^); non-selective intensity, *ρ* = –0.09 (p = 0.0003); p = 0.005, z-test; **Figure 4d**). In addition to spiking, neurons’ mean-matched Fano factor and pairwise spike-count correlations reduced with higher attentional selectivity and intensity (Fano factor, selectivity, p < 10^−315^, intensity, p < 10^−136^; pairwise correlations, selectivity, p < 10^−36^, intensity, p < 10^−6^; ANOVA; **Figure 4e-f**). A significant interaction was also detected between the selectivity and intensity on the Fano factor (p < 10^−5^, ANOVA) as well as pairwise spike-count correlations (p = 0.039, ANOVA).

**Figure 4.**
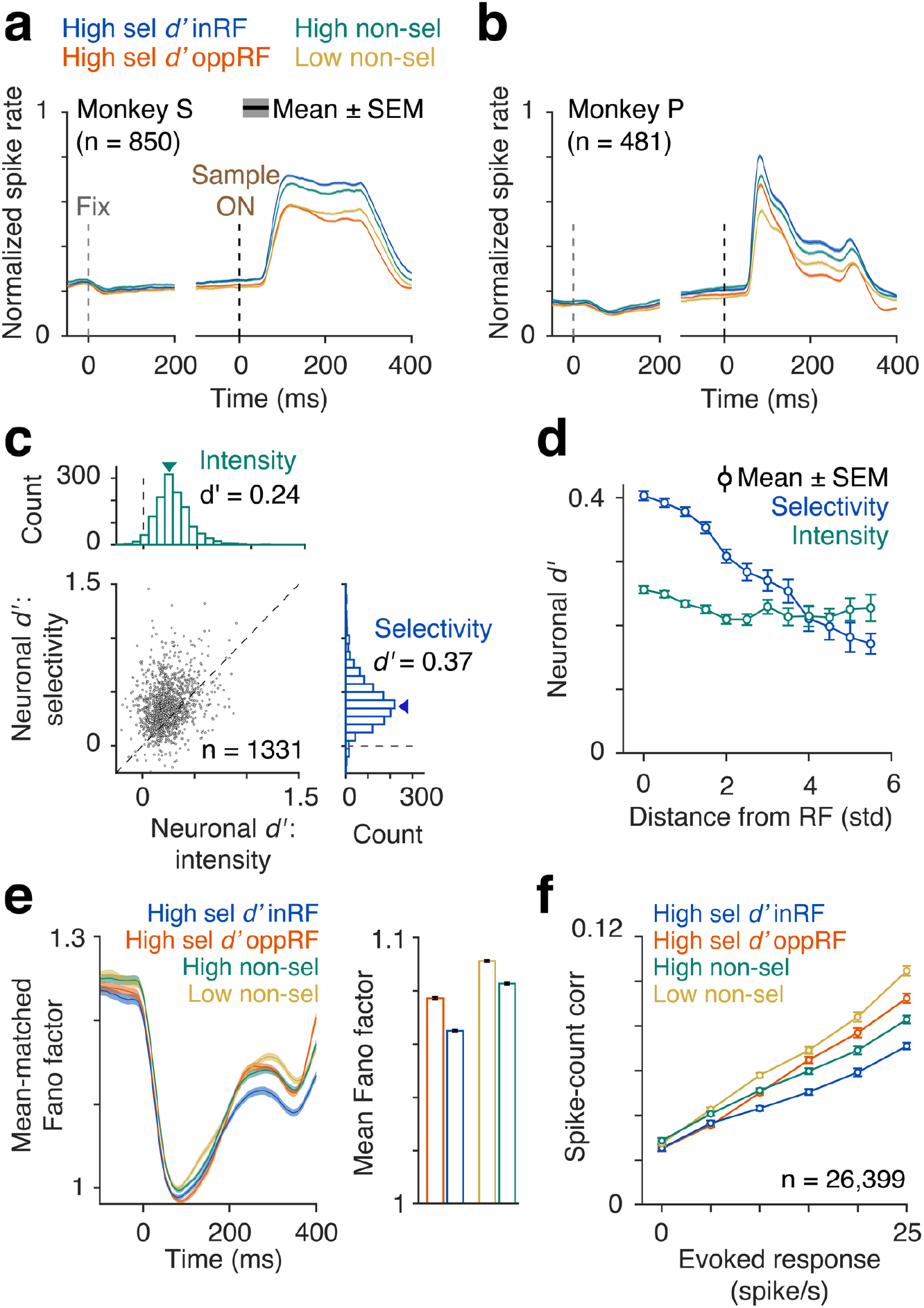
Neuronal modulation with changes in attention components controlled by differential reward size. **a-b** Population PSTHs of spike rates of correct trials in different attention conditions for monkey S (a) and monkey P (b). Single trial spike counts were binned at 2 ms, smoothed with σ = 15 ms half-Gaussian and then aligned at the onset of sample stimulus. Spike rates of each neuron were normalized to its peak response within 60 - 260 ms from sample stimulus onset (monkey S, n = 850; monkey P, n = 481). **c** Distribution of neuronal d’ for attentional selectivity and intensity of all units from both monkey S and P (n = 1331). **d** Distribution of neuronal d’ as a function of Mahalanobis distance between neurons’ RF and Gabor stimulus in units of standard deviation. **e** Left, Mean-matched Fano factor. Right, Mean-matched Fano factor averaged over 60-260 ms from the sample onset. Error bars, ±SEM. **f** Pairwise correlations between spike-counts of simultaneously recorded neurons over 60 to 260 ms from sample onset (n = 26, 399 pairs, all units) and binned according to their evoked responses (geometric mean). Error bars, ±SEM.

### Encoding of attention selectivity and intensity within single trial spike train

Spike trains of V4 neurons provide dynamic information about many task relevant variables^24^. We next measured and compared the relative contributions of attentional selectivity and intensity on the within-trial instantaneous spiking of individual V4 neurons in varying difficulty and varying reward contexts using a generalized linear encoding model^24^ (**Figure 5a**). The probability of an observed spike count within a small time window (50 ms) was modeled as an exponential function of a weighted linear combination of task variables: attentional selectivity (ratio of *d’*s at the RF location over the oppRF location), attentional intensity (radial distance from the inRF *d’*-oppRF *d’* to the origin), selectivity-by-intensity interaction, orientation of the sample stimulus inside the neuron’s RF and the direction of the eventual response saccade. The probability of a spike was constructed to follow a negative binomial distribution (**Methods**). Most neurons were well fit with this model (difficulty context, 1169/1194 (98%), p < 0.05; reward context, 1311/1331 (98%), p < 0.05; F test). **Figure 5b** shows PSTHs of observed and model fitted spike counts in different attention conditions for trials in cross-validation test data sets for an example neuron in a session with varying task difficulty. **Figure 5c** illustrates fitted model components of attention for the same example neuron as in **Figure 5b**. The effective influence of distinct attention components on spike counts are expressed as multiplicative gains (exponentiated fitted coefficients) at a representative time during the sample stimulus (150 ms). The product of these gain components results in the predicted rate for a single trial (combined gain, bottom row). For an identical increase in behavioral *d’* in the RF location, the increase in spike counts will be higher when the opposite-RF *d’* is small (blue arrow, **Figure 5c**) compared to large (green arrow, **Figure 5c**).

**Figure 5.**
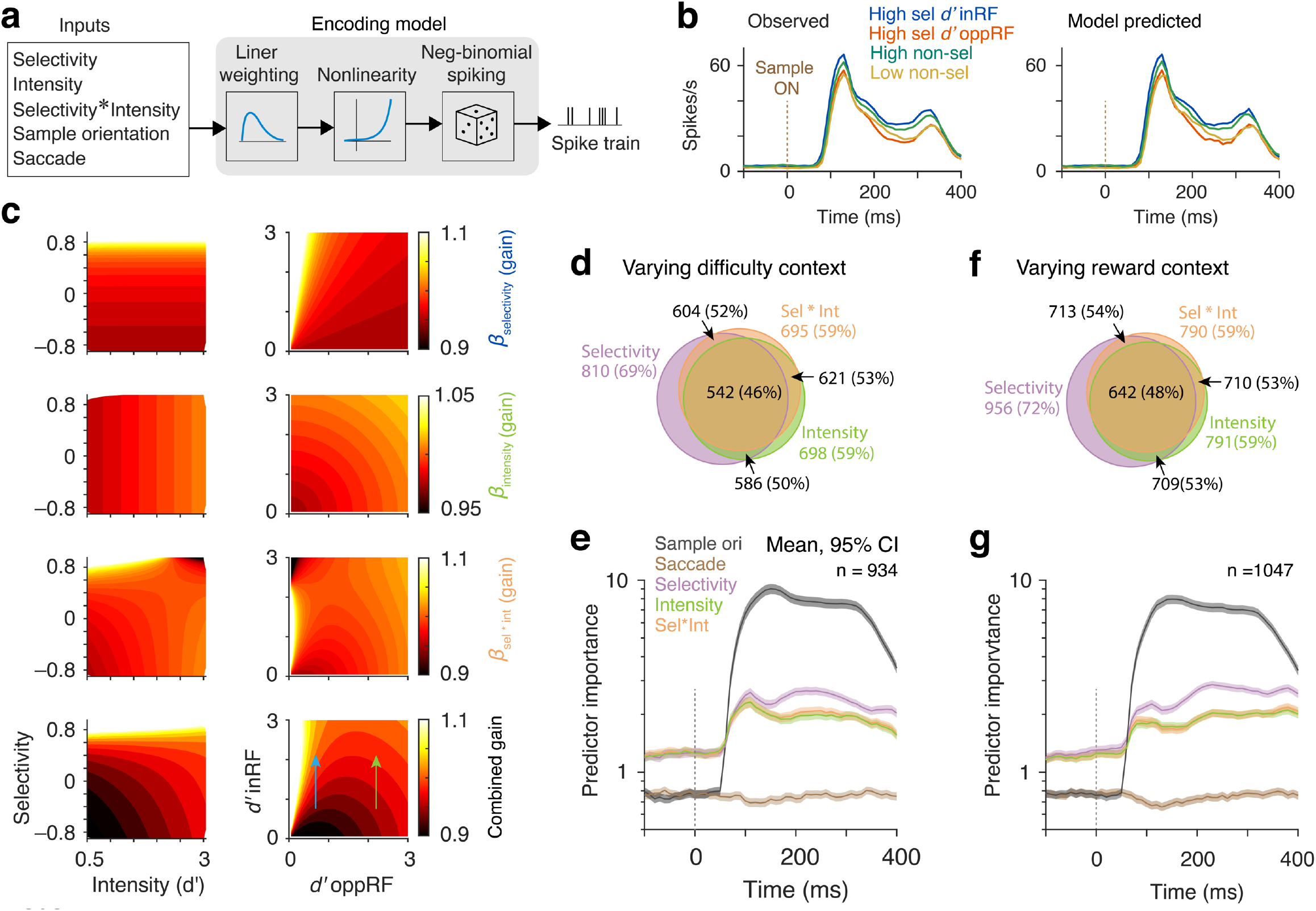
Single trial encoding of attention components in different motivation contexts. **a** Generalized linear encoding model. A neuron’s spike count over 50 ms sliding window (10 ms shift) is modeled as exponential function of linear combination of weighted (μ coefficient) experimental variables, stimulus orientation, saccade, attentional selectivity, attentional intensity and their interaction selectivity*intensity. **b** Model predicted (Left) and observed (Right) PSTHs from cross-validation test-dataset for an example neuron in varying-difficulty context. **c** Distributions of GLM fitted coefficients (exponentiated gain) for attention components at a representative time 140 ms as a function of attentional selectivity and intensity (left column), and behavioral d’s oppRF and inRF (right column) for the same example neuron in **Figure 5b**. Sel*Int, selectivity-by-intensity interaction; Combined, resultant of all 3 attention components. These components multiplicatively influence spike counts. **d-e** Varying task-difficulty context. Proportion of neurons that are modulated (p<0.05) by attentional selectivity, intensity and selectivity-by-intensity interaction estimated from the model (d). Comparing predictor importance (PI) that measures contributions of different predictor variables estimated by absolute standardized predictor coefficient values of all well fitted neurons (e). Error bars, 95% confidence intervals (bootstrap, n = 10^4^). **f-g** Varying reward context. Same as **Figure 5d** and **5e** when attention was controlled by differential reward sizes. Error bars, 95% confidence intervals.

We next ask whether attentional selectivity, intensity and their interaction were encoded by distinct set of V4 neurons and task context affects these population representations. Many units showed significant effects for several or all variables: attentional selectivity, intensity; and selectivity-by-intensity interaction (difficulty context: selectivity and intensity, 586/919 (64%), p < 10^−10^; selectivity and selectivity-by-intensity, 604/904 (67%), p < 10^−10^; intensity and selectivity-by-intensity, 621/772 (80%), p < 10^−10^; selectivity, intensity and selectivity–by–intensity, 542/934 (58%), p < 10^−10^; reward context: selectivity and intensity, 709/1038 (68%), p < 10^−10^; selectivity and selectivity-by-intensity, 713/1033 (69%), p < 10^−10^; intensity and selectivity-by-intensity, 710/871 (81%), p < 10^−10^; selectivity, intensity and selectivity–by–intensity, 642/1047 (61%). p < 10^−10^ (61%); chi^2^ test; **Figure 5d, f**). Further, these distributions were largely unchanged across the difficulty and reward contexts (p = 0.97, chi^2^ test). This mixed representation indicates multiplexed encoding of attention components by the same V4 unit.

We next compared the relative contributions of the cognitive and task variables on spike responses for individual units as measured by predictor importance (normalized magnitude of fitted coefficients, **Figure 5e, g**) (**Methods**). Following the onset of sample stimulus, stimulus orientation had a dominant contribution to the spike counts in both task contexts. This is expected because V4 neurons have robust visual responses and most are orientation selective. Attentional selectivity, intensity and selectivity-by-intensity interaction remained strong predictors of spike trains from the start of the trial, and increased immediate after the stimulus onset. Saccade direction contributed negligibly to V4 activity. Contributions of attention components were indifferent to the task contexts. Both animals showed similar results (**Supplementary Figure S3, S5**). Collectively, these results suggest that individual V4 neurons independently carry multiplexed information about selective attention and attention intensity in single trial spike trains relative to other sensory and task variable. These two attention components are integrated independent of the way the animal is motivated to allocate their attention.

### A normalization model of attention can account for the interactions between attentional selectivity and intensity

The decrease in V4 responses with the increase in behavioral *d’* in the opposite hemifield (blue to green, yellow to orange, **Figure 2d, 3c**) might seem unexpected, both because behavioral performance at the receptive field location does not change and because overall behavioral performance is better when the animal allocates high attention to both locations. This reduced rate of firing can be understood in the context of spike response normalization. To gain a mechanistic understanding of the observed interactions between attentional selectivity and non-selective intensity seen in V4 responses, we tested whether a simple extension of a sensory normalization model^25^ with spatially distributed behavioral *d’* can account for these effects on spike responses. Mean stimulus evoked spike counts were expressed as: *r* = (d’_in_*E_in,G_ + d’_opp_*E_opp,G_)/(d’_in_*S_in,G_ + d’_opp_*S_opp,G_ + *σ*); where *d’*_*i*_ represents behavioral *d’* at location *i* in or opposite the RF hemifield, E_i,G_ and S_i,G_ represent excitation and suppression at location *i* due to the Gabor stimulus (G = 1) or the background (G = 0), and σ is baseline suppression (**Figure 6a** and **Methods**). Model parameters E_i,G_, S_i,G_ and α were fit for each unit with the trial-averaged spike counts over 200 ms during the pre-stimulus fixation, sample and test interval periods on training datasets. Performance of the full model (*model with d’*) was measured on the 4-fold cross-validation test datasets subsampled across different stimulus configurations and attention conditions, and compared with two alternate models that lacked any behavioral *d’* factors: *model without d’*, and *model without d’ and background display* (**Methods**)

**Figure 6.**
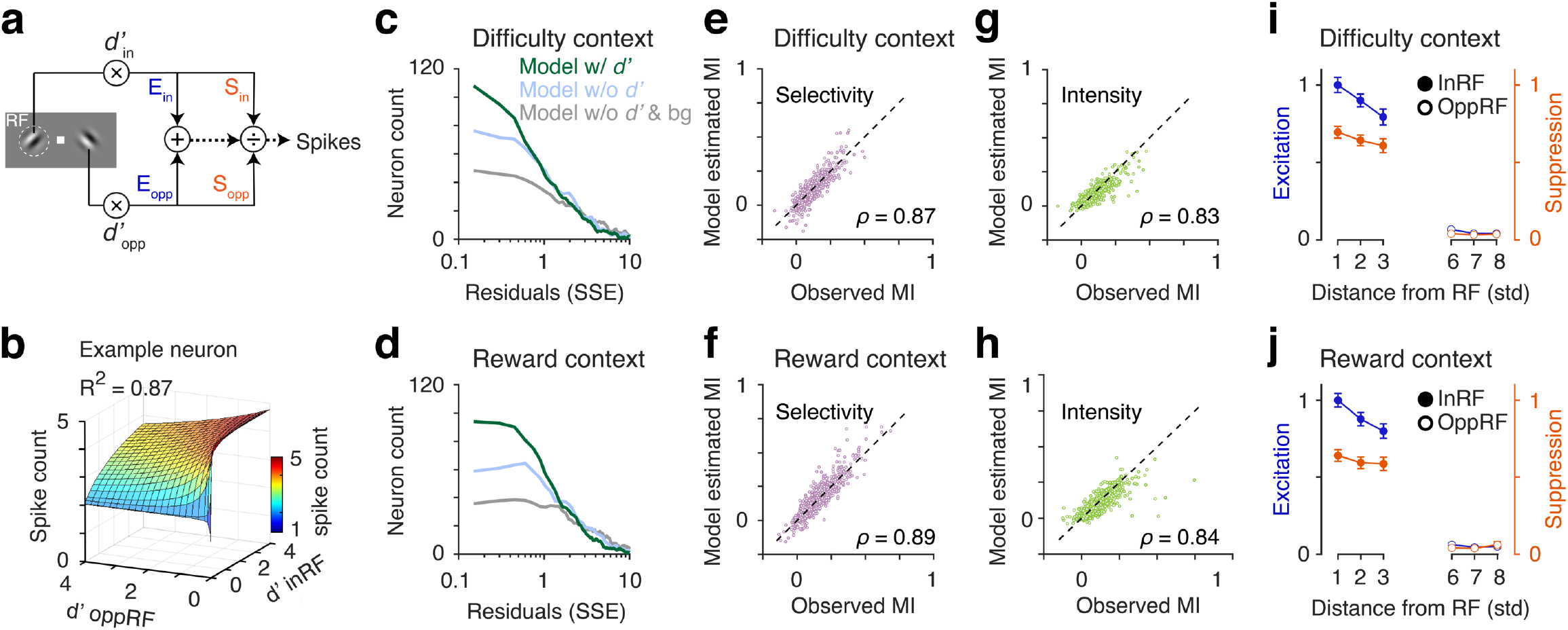
Normalization model of attention can account for attentional intensity effects on spike counts. **a** Normalization model of attention. Spike count, r = (d’_in_*E_in,G_ + d’_opp_*E_opp,G_)/(d’_in_*S_in,G_+ d’_opp_*S_opp,G_ + α). Where, d’_i_ is the behavioral d’ at location i (inRF or oppRF). E_i,G_, S_i,G_ are respectively excitation and suppression at location i due to either the Gabor stimulus (G = 1) or the background (G = 0). α is a constant. Spikes counts were fitted with 3 different models (“Methods”). Model w/ d’: contains behavioral d’ values in two stimulus locations. Additionally, there are excitation and suppression terms for background display. Model w/o d’: same as the previous model except without the d’ terms (d’_i_ = 1). Model w/o d’ & bg: Same as the previous model, except without any d’s and excitation/suppression parameter due to the background (E_i,G=0_= 0, S_i,G=0_ = 0). **b** Fit of an example neuron with the normalization model w/ d’. Surface plot, fitted spike counts. **c-d** Quality of fit (SSE) for all recorded units in two behavioral contexts (difficulty and reward) fitted with the three different normalization models. Spike-counts of the most of the neurons were well fit with the normalization model w/ d’ compared to w/o d’ models. **e, g** Comparing spike-count modulations (modulation index, MI) with attentional selectivity (e) and intensity (g) between observed and model fits for the cross-validation test datasets across difficulty context sessions. MIs from observed and fitted spike counts are highly correlated (Spearman correlation coefficient for selectivity, ρ = 0.87, p < 10^−100^; for intensity, ρ = 0.83, p < 10^−100^). **f, h** Same as in (e) and (g) for reward context sessions (Spearman correlation coefficient for selectivity, ρ = 0.89, p < 10^−100^; for intensity, ρ = 0.84, p < 10^−100^). **i-j** Population averaged model fitted excitation (E) and suppression (S) parameters from the normalization model w/ d’ across all units binned according to the distance (Mahalanobis) of unit’s RF location from the Gabor stimulus (same or opposite of the RF hemifield).

The surface plot in **Figure 6b** shows mean spike counts fitted with the normalization model (*model with d’*) as a function of behavioral *d’*s in two hemifields for an example unit in a difficulty context session. Most units in the two task contexts were better fit with the attention model of normalization compared to alternate *d’*-independent normalization models (difficulty session: *model with d’*, 910/1194 (76%), *model without d’*, 626/1194 (52%), *model without d’ and background display*, 479/1194 (40%), reward session: *model with d’*, 1006/1331 (76%), *model without d’*, 614/1331 (46%), *model without d’ and background display*, 486/1331 (37%); variance explained >80%; **Figure 6c-d**). At the population level, the quality of normalization model fits of spike responses in the two task contexts did not differ (p = 0.23, chi^2^ test). The normalization model with *d’* captured multiple features of observed spike counts across different stimulus and attention conditions, including relative changes in spike counts across different and attention conditions, and neuronal modulation indices for attentional selectivity and intensity (**Supplementary Figure S7, Figure 6e-h**). Population correlation between the observed and normalization model estimated spike count modulation indices for attentional selectivity and intensity were strongly correlated across the task contexts (difficulty context: attentional selectivity, *ρ* = 0.87, p < 10^−10^, attentional intensity, *ρ* = 0.83, p < 10^−10^; reward context: attentional selectivity, *ρ* = 0.89, p < 10^−10^, attentional intensity, *ρ* = 0.84, p < 10^−10^; Spearman correlation coefficient; **Figure 6e-h**). Further, the model fitted excitatory and suppressive stimulus drives of recorded neurons decreased with the proximity of the neuron’s RF and the stimulus irrespective of task context that motivated animals to attend (**Figure 6i-j**). Together, these results show that a simple normalization model captures the effects of attentional engagement at a distant site on the spike response in the RF location regardless of the stimulus or attentional context of that RF response.

## DISCUSSION

We isolated the contributions of attentional selectivity and non-selective intensity to the activity of individual V4 neurons while precisely and independently controlling monkeys’ behavioral *d’* at the RF location and a distant location in the opposite hemifield. Changes in either attentional selectivity or intensity independently are associated with overall increases in V4 spike rates and decreases in V4 spiking variability and pairwise spike count correlations. Further, spike rates were reduced when behavioral d’ increased at a distant location in opposite hemifield. A spatially tuned response normalization model explained all these changes in spike rate across attention conditions and task contexts. Finally, single trial encoding of attentional selectivity, intensity and their interaction in V4 neurons were found to be independent of the way the subject is motivated to regulate its attention.

A functional role for the spiking of V4 neurons is supported by a correlation with enhanced sensory processing of an attended stimulus at the RF location, as well as impaired behavioral detection in subjects with V4 lesions. V4 lesions in macaques and humans impair attentional performance by making it difficult for the subjects to exclude irrelevant distractors^26,27^. However, our results reveal a mismatch between V4 spiking and behavioral performance: lower spike rates for the same behavioral performance when monkey’s attention strategy shifts from spatially selective to non-selective high intensity at the RF location (**Figures 2b, 4a**). This raises questions about the relationship between neuronal signals in V4 and a subject’s perceptions and performance. Other studies also found a dissociation between behavioral performance and activity in cortical visual areas, including V4^5,24,28^. That work showed dissimilar dynamics of top-down attentional control signals, sensory modulation and executive action. Specifically, when attention shifted, behavioral changes lagged changes in the spike rates of sensory neurons by seconds to minutes, breaking the link between spikes and performance^24^. Our results show that a mismatch between V4 activity and behavior can also go beyond lags: when animals spread their field of attention beyond the RF, V4 spike rates go down even though the animals maintained the same behavioral *d’*. While a normalization mechanism can explain the reduced spiking (**Figure 6**) it does not address this disconnect with behavior. Changes in pupil diameter suggest that the animal in fact increased its total effort at both sites to maintain performance in the high effort condition (**Figures 1f, 3e**), making the reduced spike rates even less expected. Nevertheless, there is little reason to believe that behavioral performance should be uniquely determined by the strength or quality of sensory signals in any one brain region. A primate brain doing an attentionally-demanding task depends on contributions from many structures throughout the neuraxis. Even if perceptual stages perform perfectly, overall behavioral performance can be affected by distractions and lapses associated with activity in other brain stations. The somewhat reduced spike rates in V4 that occur with high attention to both hemifields might in fact lower performance, but be counterbalanced by attention-related changes in other structures that enhance performance by reducing errors related to factors like distractions or motor error.

In the current experiments, monkeys were motivated to direct their spatial attention by expectation of either larger rewards or higher task demands associated with two locations. The invariance of neuronal modulations in V4 with these two distinct motivational factors suggest that these effects depend on a common top-down cognitive control and do not represent reinforcement signals to any appreciable extent. This notion is further supported by previous reports of weak evidence of encoding of reward information by single trial spike counts of V4 neurons across different attention states and during state transitions^24^ (but see^5^). Motivation plays a crucial role in influencing attention control, giving priority to the most appropriate goal among multiple competing targets. Both humans and animals can be motivated to execute actions either for intrinsic pleasure or for satisfying some basic needs such as hunger, thirst, etc. Several brain structures within the frontoparietal network are modulated by motivationally salient signals such as errors, rewards and penalties^29^. Although various forms of motivations (intrinsic and extrinsic) can have different origins, they have common nodal points in the striatum and prefrontal cortex that receive dopaminergic afferents that play a crucial role in reward learning^30^. Consistent with this, previous evidence points to common cortico-limbic neural pathways that are activated by either changes in expectation of reward or changes in task difficulty^31^.

Our results show a close relationship between attentional intensity and “effort”. Previous work has associated cognitive effort exclusively with changes in task difficulty, and viewed it as a specific type of “arousal” that is distinguishable from other forms of arousal elicited by exogeneous factors such as stress, novel stimuli and drugs^16^ (but see^32^). The manifestation of attentional intensity on regulating visual sensory processing of V4 neurons in both of our task contexts instead supports identifying effort as an intensive aspect of attention^16^. It is possible that the component of bottom-up stimuli driven arousal that affects performance might map well onto the neuronal modulation associated with attentional intensity or effort. Future experiments with precise and independent control of these cognitive components in simultaneous tasks might identify their precise relationships and how subjective experience of effort relates to attentional intensity. Other important questions to be addressed concern the circuit, cellular and molecular mechanisms that mediate attentional intensity. These could involve activation of diverse neuromodulatory systems such as norepinephrine, acetylcholine, serotonin^6,33-37^.

V4 spike responses for varying attentional selectivity and intensity were well explained by an extension of normalization model of visual responses^38^ with spatially tuned attentional gain factors represented by behavioral *d’*. A similar normalization model with uniform attentional effects on excitation and surround suppression have been previously used in explaining neuronal modulations in V4 with spatially selective attention^25^. Previous normalization models considered attentional effects on stimulus-induced excitation and surround suppression. To explain a modest but non-zero neuronal modulation during the fixation period in absence of any stimulus, our normalization model included an attention effect on the background display similar to the visual stimulus. This is consistent with the evidence that attention acts as a constant gain factor^3^. Similar to reward expectations, increased task difficulty associated with spatial attention increases visual excitation^13^ and response suppression^20^ of V4 neurons. Further, response suppression with high task load is considered to serve as a mechanism for reducing peripheral interference and improving signal detection^39^. These reports are inconsistent with the correlated decrease in model estimated excitation and suppression with an increase in the proximity of neuron’s RF and attended stimulus across task difficulty and reward expectation contexts. Together, this normalization model of attention provides a canonical neuronal computation to explain how distributed spatial attention influences neuronal responses. Future experiments are required to examine how other forms of attention such as feature-based or bottom-up attention act on normalization mechanisms across different visual areas that are responsive to attentional modulation.

Taken together, our results provide new experimental evidence revealing how attentional selectivity and non-selective intensity interact and modulate sensory processing in visual cortex in reference to behavioral performance. Moreover, our study identified a computational mechanism of normalization through which spatially distributed attentional performances interact.

## METHODS

### Subjects and surgery

Two adult male rhesus monkeys (*Macaca mulatta*, 13 and 9 kg) were implanted with a titanium head post using aseptic surgical techniques before training began. After the completion of behavioral training (3 to 5 months), we implanted a 10×10 array microelectrodes with 400 μm spacing (Blackrock Microsystems) into dorsal visual area V4 of one hemisphere, between lunate and superior temporal sulci. The same two monkeys were used in a previous study that included some of the same neuronal responses, but described different findings^24^.

### Behavioral task

During training and neurophysiological recording, the monkey sat in a primate chair facing a calibrated CRT display (1024 x 768 pixels, 100 Hz refresh rate) at 57 cm viewing distance inside a darkened room. Binocular eye position and pupil area were recorded at 500 Hz using an infrared camera (Eyelink 1000, SR Research). Trials started once the animal fixated within 1.5° of a central white spot (0.1^°^ square) presented on a mid-level gray background (**Figure 1a**). The animal had to maintain fixation until its response at the end of the trial. After a fixation period of 400-800 ms, two achromatic Gabor sample stimuli appeared for 200 ms, one in each visual hemifield. After a variable delay of 200-300 ms, a Gabor test stimulus (test 1) appeared for 200 ms at one of the two target locations, randomly selected with equal probability. The test stimulus was identical to the preceding sample stimulus, except potentially its orientation. On half of the trials, the test 1 stimulus had a different orientation (nonmatch trial) and the monkey had to make a saccade to that target to receive an apple juice reward. On the remaining half of the trials, the test 1 stimulus had the same orientation as the corresponding sample stimulus (match trial), and the monkey had to maintain fixation until a second test stimulus with a different orientation (test 2, 200 ms) appeared in the same location after an additional delay of 200-300 ms. The monkey then had to saccade to that target to get a reward. Inter-trial intervals varied from 2-3 s. Stimuli were presented always in the lower hemifields at 2°-4° eccentricity. Gabors were static and odd-symmetric with the same average luminance as the background. Spatial frequency, size and base orientation of Gabor stimuli were optimized for one neuron recorded each day, and remained unchanged throughout each session (left, azimuth –2.5° to –4.5°, elevation –0.5° to –4.0°, sigma, 0.35° to 0.70°, spatial frequency 0.6 to 3.5 cycles/°; right, azimuth 1.8° to 5.5°, elevation –0.5° to –4.0°, sigma, 0.25° to 0.58°, spatial frequency 0.7 to 3.0 cycles/°). On every trial, the orientation of the sample stimuli randomly took one of two values (independently), base orientation or orthogonal. Stimulus parameters and orientation changes remained fixed within a session and varied across sessions. Orientation changes differed between blocks when task difficulty was manipulated, but every block contained probe trials that had the same orientation change throughout a session (24°-40° for monkey S, 20°-40° for monkey P). Reward sizes for hits (correct response in nonmatch trial) and CRs (correct rejections in match trial) were adjusted by < 10% as needed to encourage the animal to maintain a behavioral criterion close to zero. Behavioral task was controlled using custom-written software (https://github.com/MaunsellLab/Lablib-Public-05-July-2016.git).

#### Behavioral task contexts

Animals were motivated to allocate their spatial attention using two different task contexts, varying task difficulty or varying reward size. In alternate sessions, animal’s spatial distribution of behavioral *d’* at two locations in opposite hemifields was controlled by either of the task contexts. In task-demand context, selective attention and non-selective attentional intensity were controlled over interleaved blocks of trials (160-440 trials/block) by changing relative task difficulty at the two locations in opposite hemifields (**Figure 1c**). Orientation change randomly took one of two values, probe orientation change (∼30% of the trials) or contextual orientation change (∼70% of the trials). Contextual orientation change was small for difficult task (high task-demand) and large for easy task (low task demand) compared to the probe orientation change. A high behavioral *d’* (selective attention) at location 1 relative to the location 2 was achieved by making the task difficulty high at the location 1 (ΔΘ_context_, 17°-20° for monkey S, 8°-22° for monkey P) and easy at the location 2 (ΔΘ_context_, 80°-90° for monkey S, 80°-90° for monkey P). A high non-selective behavioral *d’* (high non-selective attention intensity) was obtained by making the task difficulty high at both the locations (ΔΘ_context_, 15°-18° for monkey S, 6°-22° for monkey P). A low non-selective behavioral *d’* (low non-selective attention intensity) was obtained by making the task difficulty easy at both the locations (ΔΘ_context_, 80°-90° for monkey S, 80°-90° for monkey P) (**Figure 1b**). Reward values for correct behavior responses were always the same across blocks on both sides and fixed.

In the differential reward context, selective attention and non-selective attention intensity were controlled over interleaved blocks of trials (120-220 trials/block) by changing reward size at the two locations (**Figure 3b**). There was only a single orientation change that remained fixed throughout the experiment session. A high selective behavioral *d’* (selective attention) at location 1 relative to the location 2 was achieved by delivering high rewards at the location 1 compared to the location 2. High non-selective behavioral *d’* (high non-selective attention intensity) was controlled by delivering high rewards for correct responses at both locations. Similarly, a low non-selective behavioral d’ (low non-selective attention intensity) was controlled by giving low rewards for the correct responses at both locations. Animals were encouraged to maintain a behavioral target/non-target criterion close to zero by small adjustment of trial-by-trial reward ratio for hits and correct rejections (**Supplementary Figure S1**).

### Electrophysiological Recording and Data Collection

Extracellular neuronal signals from the chronically implanted multielectrode array were amplified, bandpass filtered (250–7,500 Hz) and sampled at 30 kHz using a data acquisition system (Cerebus, Blackrock Microsystems). We simultaneously recorded from multiple single units as well as multiunits over 42 differential task-demand sessions (714 units and 20 sessions for monkey S; 480 units and 22 sessions for monkey P) and 36 differential reward sessions (850 units and 20 sessions for monkey S; 481 units and 16 sessions for monkey P). At the start of each experimental session we mapped RFs and stimulus preferences of neurons while the animal fixated. These RFs were used to optimize the stimulus parameters. Spikes from each electrode were sorted offline (Offline-Sorter, Plexon Inc.) by manually well-defining cluster boundaries using principal component analysis as well as waveform features. Well isolated clusters were classified as single units from multiunits based on the isolation quality of unit clusters. The degree to which unit clusters were separated in 2D spaces of waveforms features (first three principal components, peak, valley, energy) was measured by Multivariate Analysis of Variance (MANOVA) F statistic using Plexon offline Sorter (Plexon Inc.). A unit cluster of MANOVA p-value of < 0.05 was considered as single unit which indicates that the unit cluster has a statistically different location in 2D space, and that the cluster is statistically well separated.

### Data Analysis

#### Behavioral Sensitivity (d’), criterion

All completed trials in the reward context and all probe orientation trials in task-demand context were included in our analysis. Behavioral sensitivity (*d’*) and criterion (*c*) at a spatial location were measured from hit rates within nonmatch trials and FA rates within match trials as:

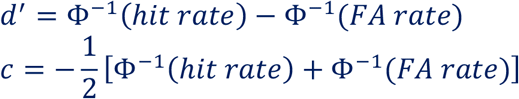

where Φ^−1^ is inverse normal cumulative distribution function. We measured average *d’* and *c* within a session across all trials across blocks separately for four different attention conditions.

#### Index of attentional selectivity and intensity

Attention selectivity was measured by the relative value of the behavioral *d’* inside the RF location with respect to opposite RF location. The measure mapped directly onto polar angle in *d’*-space (**Figure 1d, 3c**) and was normalized to a range from -1 (inside RF *d’* = 0) to 1 (opposite RF hemifield *d’* = 0):

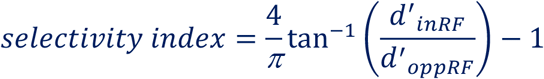

Where, 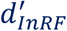 and 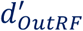 are the sensitivities in the two hemifields, inside and outside the recorded neurons’ RFs. Attentional intensity represented the absolute value of total behavioral d’ (distance from the origin in d’ space):

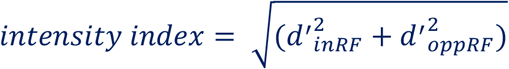

#### Pupil area

All pupil area measurements were measured binocularly at 500 Hz while monkeys maintained fixation in absence of a luminosity change using infrared camera (EyeLink 1000, SR Research). Raw pupil areas were z-scored for each session and each eye separately. Mean pupil area was measured by averaging the z-scored pupil area during the 400 ms after sample appearance.

#### Neuronal response modulation

Only neurons with an average spike rate 60-260 ms after sample stimulus onset that was significantly (p < 0.05) greater than the rate 0 to 250 ms before sample onset were used in the analysis. To construct peri-stimulus time histograms (PSTHs) for figures, spike trains were aligned to sample stimuli onset and averaged across trials and smoothed with a half Gaussian kernel (rightward tail, SD 15 ms). A spike rate modulation (**Figure 2c, 2d, 4c and 4d**) was quantified by neuronal *d’* as:

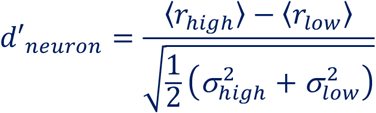

Where <*r*_*x*_> and σ_x_ are the average and SD of the spike counts within 60 to 260 ms from sample stimuli onset. Neuronal *d’*s were calculated for each unit separately for attentional selectivity and intensity. Neuronal modulation index (MI, **Figure 6e-h**) was measures as:

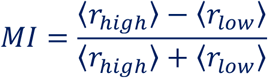

#### Proximity between neuron’s receptive field (RF) and sample stimulus

Proximity between neuron’s receptive field (RF) and sample stimulus was estimated by the Mahalanobis distance (standardized distance, **Figure 2d** and **4d**). For each neuron, spatial RF was measured and fit using a bivariate Gaussian. We then calculated the Mahalanobis distance between probability densities of spatial RF and the Gaussian contrast profile of the Gabor stimulus.

#### Fano factor

Mean-matched Fano factor (**Figure 2e** and **4e**) was measured using spike counts over 100 ms sliding windows in 2 ms steps for each neuron according to procedures described previously^24,40^. Then the variance and mean across trial was computed at every time bin. The greatest common distribution of means across neurons, attentional intensities and time bins were measured. In order to match the mean distribution to the common mean distribution, a different subset of neurons was randomly chosen (50 times) at every time bin and the average Fano factor was computed (ratio of the variance to the mean).

#### Spike-count correlations

Pearson correlation coefficients were computed for pairs of simultaneously recorded units on spike-counts over 200 ms (60-260 ms from sample stimuli onset), defined as the covariance of spike counts normalized by the variances of individual neurons:

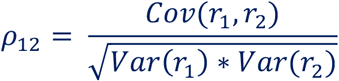

Where r_1_ and r_2_ are spike counts of neuron 1 and neuron 2 across trials. Pairwise spike-count correlations were binned according to the geometric mean of the evoked responses of the two neurons in 5 Hz intervals. Evoked response was computed and subtracting by the trial-averaged baseline spike rate (–200 to 0 ms from sample onset) from the trial-averaged spike rate during the sample (60-260 ms from sample onset) (**Figure 2f** and **4f**).

#### Generalized linear encoding model

Generalized linear model (GLM) regression was used to estimate the encoding of different attention components and task variables in single trial spike trains. Spike counts (*r*) over 50 ms bins with 10 ms shift in single trials were modeled to follow a negative binomial distribution. The negative binomial distribution is well suited for the purpose, as spike count variances of cortical neurons are most often equal to or greater than their means^24,40,41^. The details of the model implementation were described in an earlier study^24^. Briefly, expected value of spike count at each time bin according to the GLM was represented as:

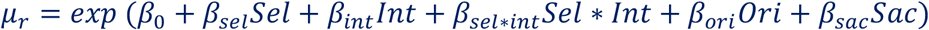

Where, *β*_*i*_ is the coefficient for the predictor variable *i*; Sel, session averaged attentional selectivity, ratio of the behavioral *d’* at the RF location to the *d’* at the opposite hemifield location; Int, session averaged attentional intensity, distance from the origin in *d’* space; Sel*Int, interaction between selectivity and intensity; Ori, orientation of sample stimulus inside neuron’s RF; Sac, saccade choice (1 for saccade towards the RF, –1 for saccade opposite to the RF and 0 for saccade withheld). For the reward context dataset, we also used another alternate GLM containing an additional predictor variable, reward history.

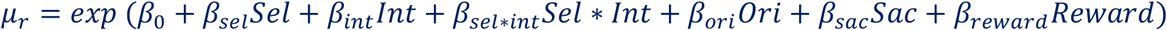

Here, reward represents average of reward values on the 3 immediately preceding trials. GLM was implemented in Matlab separately for each neuron. In order to compare different predictor coefficients, predictor variables were converted to z-scored values and fitted with GLMs to obtain standardized beta-coefficients. Goodness of fit for a given GLM was measured by residual deviance, pseudo R-squared value (Cragg & Uhler’s method) and F-statistics compared to a null model.

Predictive performance of the GLMs was measured by cross validation. Observations in each neuron’s dataset were split at random into 10 partitions. GLM fit was done on 9 training partitions and the remaining partition was used for cross-validation. This cross-validation error for each neuron was measured by:

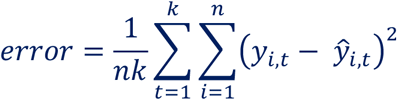

Where *n* is the number of cross-validation trials; *k* is the number of time bins; *y*_*i,t*_ and *ŷ*_*i,t*_ are respectively recorded and GLM estimated spike count at *t*^*th*^ time bin in *i*^*th*^ trial. The quality of cross validation for neuron populations was measured by the Spearman correlation coefficients between the observed and GLM fit spike counts.

Relative importance of each predictor variable was measured by predictor importance (PI) expressed as the absolute z-statistic of each fitted predictor coefficient:

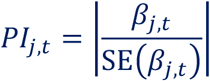

Where *j* = 1, 2, 3,…, *m*, predictor variables; *t* is time bin. A neuron with predictor coefficient different from zero (p < 0.05; t-test) during the sample stimuli presentation (60 - 260 ms from sample on) was classified as sensitive to that predictor. Because of the exponential nonlinearity in the GLM, exponentiated fitted predictor coefficients were used to illustrate how attentional selectivity, intensity and their interaction (selectivity-by-intensity) influence on spike rates as a function of attentional selectivity and intensity (left, **Figure 5c**). as well as a function of behavioral *d’*s at the RF location and opposite hemifield location (right, **Figure 5c**). These components are combined multiplicatively to drive instantaneous spike rates of individual units.

#### Normalization model

Trial averaged spike counts over 200 ms across all attention conditions and stimulus configurations were fit with three different normalization models. Two were simple stimulus normalization models without any spatial tuning (*model without d’ & background display* and *model without d’*) and the third model was an extension of spatially tuned normalization model (*model with d’*). All the 3 models were fit with 9 non-negative parameters and used the same set of 27 training data points out of 36 using nonlinear least-squares solver (MATLAB lsqnonlin.m, MathWorks). The quality of fit was measured by residuals using 9 cross-validation test data points (4-fold cross-validation). The training dataset consisted of mean spike counts during 3 pre-sample (–200 to 0 ms), 12 sample stimuli (60 -200 ms) across two Gabor orientations and four attention conditions, 6 test 1 stimulus (60 - 200 ms) on left hemifield and 6 test 1 stimulus on right hemifield across all four attention conditions and two Gabor orientations. Cross-validation (fourfold) test dataset contained spike counts during 1 pre-sample, 4 during the sample stimuli, 2 during test 1 stimulus inside the RF location and 2 during test 1 stimulus in the opposite RF location across all attention conditions and Gabor stimulus configurations.

##### Model without *d’* and without background display

Mean spike counts during the sample stimuli period according to the normalization *model without d’ & without background display* is described as:

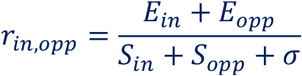

Where fit parameters *E*_*i*_, *S*_*i*_ respectively represents excitation, suppression drives due to a Gabor stimulus at *i*^*th*^ location (inRF or oppRF) and *σ* represents a constant baseline suppression. For every *E*_*i*_ (or *S*_*i*_), there are two parameters, *E*_*i,base*_ and *E*_*i,base+90*_ (or *S*_*i,base*_ and *S*_*i,base+90*_) associated with each of the two Gabor orientations (base and base + 90°). The contrast term was 1 for the Gabor stimulus and 0 for no-stimulus. Thus, the mean spike counts during the inRF and oppRF test 1 presentations respectively are:

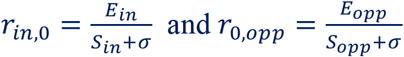

The mean of spike counts during the pre-sample period was zero.

##### Model without *d’*

According to the normalization model w/o *d’*, mean spike counts during the sample, test 1 and pre-sample periods are respectively described as:

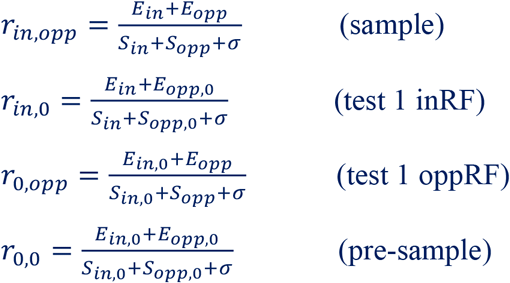

Where, *E*_*i*_ (or *S*_*i*_) could be either *E*_*i,base*_ and *E*_*i,base+90*_ (or *S*_*i,base*_ and *S*_*i,base+90*_), excitatory (or suppressive) stimulus drives at *i*^*th*^ location (inRF or oppRF) due to the two Gabor orientations; *E*_*i,0*_ (or *S*_*i,0*_) is the excitatory (or suppressive) drive due to the background display in absence of Gabor stimulus. Only one common parameter was used for excitatory (or suppressive) drives at the opposite RF location for both Gabor stimulus as well as background display, i.e., *E*_*opp,base*_ = *E*_*opp,base+90*_ = *E*_*opp,0*_ and *S*_*opp,base*_ = *S*_*opp,base+90*_ = *S*_*opp,0*_. Total 9 parameters were fit with the model.

##### Spatially tuned normalization model (*model with d’*)

According to the normalization model with *d’*, mean spike counts during the sample, test 1 and pre-sample periods are respectively described as:

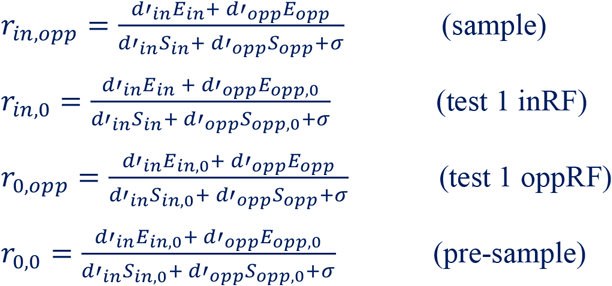

Where, *d’*_*in*_ and *d’*_*opp*_ are respectively behavioral *d’* at the RF and opposite RF locations. The fit parameters are same as in *model without d’*. In **Figures 6i-j**, excitatory (or suppressive) drives across stimulus types were averaged at the RF location for each neuron and the then the neurons sorted according to the distance

### Statistical analysis

Unless otherwise specified, we used paired t-test and multifactor ANOVA for comparing normally distributed datasets. Normality was checked using a Kruskal-Wallis test.

## Acknowledgements

We thank Jackson J. Cone, Chery Cherian for helpful discussion and/or comments on the manuscript.

