## Supplementary Information for "Neuronal correlates of attentional selectivity and intensity in visual area V4 are invariant of motivational context"

#### Supplementary Tables

**Supplementary Table T1. Behavioral d’ controlled by differential task difficulty**

| **Monkey S** (N = 20) | $d'_{inRF}$ (Mean ± SEM) | $d'_{oppRF}$ (Mean ± SEM) | Selectivity index (Mean ± SEM) | Intensity (Mean ± SEM) |
|---|---|---|---|---|
| Low selective inRF | 0.96 ± 0.04 | 2.79 ± 0.07 | –0.58 ± 0.01 | 2.96 ± 0.07 |
| High selective inRF | 2.64 ± 0.06 | 1.03 ± 0.03 | 0.52 ± 0.01 | 2.84 ± 0.06 |
| Non-selective low intensity | 0.98 ± 0.04 | 1.07 ± 0.03 | –0.05 ± 0.02 | 1.46 ± 0.04 |
| Non-selective high intensity | 2.63 ± 0.06 | 2.78 ± 0.06 | –0.04 ± 0.01 | 3.84 ± 0.08 |

| **Monkey P** (N = 22) | $d'_{inRF}$ (Mean ± SEM) | $d'_{oppRF}$ (Mean ± SEM) | Selectivity index (Mean ± SEM) | Intensity (Mean ± SEM) |
|---|---|---|---|---|
| Low selective inRF | 1.28 ± 0.06 | 2.53 ± 0.08 | –0.41 ± 0.02 | 2.84 ± 0.09 |
| High selective inRF | 2.85 ± 0.08 | 1.29 ± 0.05 | 0.46 ± 0.01 | 3.14 ± 0.09 |
| Non-selective low intensity | 1.20 ± 0.05 | 1.20 ± 0.06 | 0.01 ± 0.02 | 1.71 ± 0.07 |
| Non-selective high intensity | 2.71 ± 0.08 | 2.62 ± 0.1 | 0.02 ± 0.02 | 3.78 ± 0.11 |

**Supplementary Table T2. Behavioral d’ controlled by differential reward size**

| **Monkey S** (N = 20) | $d'_{inRF}$ (Mean ± SEM) | $d'_{oppRF}$ (Mean ± SEM) | Selectivity index (Mean ± SEM) | Intensity (Mean ± SEM) |
|---|---|---|---|---|
| Low selective inRF | 0.79 ± 0.05 | 2.73 ± 0.07 | –0.64 ± 0.02 | 2.84 ± 0.08 |
| High selective inRF | 2.61 ± 0.08 | 0.98 ± 0.06 | 0.54 ± 0.02 | 2.80 ± 0.08 |
| Non-selective low intensity | 0.99 ± 0.03 | 1.05 ± 0.04 | –0.04 ± 0.02 | 1.45 ± 0.05 |
| Non-selective high intensity | 2.53 ± 0.08 | 2.72 ± 0.07 | –0.05 ± 0.01 | 3.72 ± 0.1 |

| **Monkey P** (N = 16) | $d'_{inRF}$ (Mean ± SEM) | $d'_{oppRF}$ (Mean ± SEM) | Selectivity index (Mean ± SEM) | Intensity (Mean ± SEM) |
|---|---|---|---|---|
| Low selective inRF | 1.10 ± 0.05 | 2.51 ± 0.05 | –0.47 ± 0.02 | 2.75 ± 0.05 |
| High selective inRF | 2.66 ± 0.05 | 1.15 ± 0.05 | 0.48 ± 0.02 | 2.91 ± 0.06 |
| Non-selective low intensity | 1.05 ± 0.06 | 1.11 ± 0.06 | –0.04 ± 0.03 | 1.53 ± 0.07 |
| Non-selective high intensity | 2.68 ± 0.07 | 2.54 ± 0.08 | 0.04 ± 0.02 | 3.70 ± 0.09 |

### Supplementary Figures

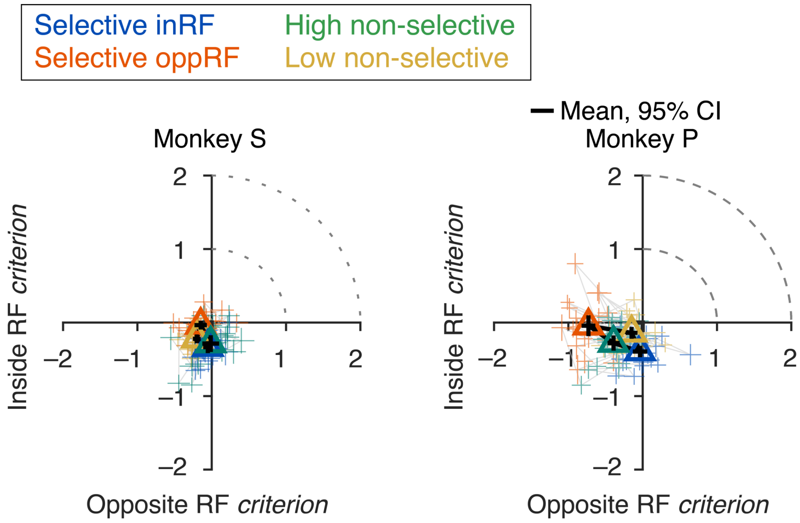


**Supplementary Figure S1. Behavioral criterions associated with attentional selectivity and intensity controlled by task difficulty.** Attention operating characteristic (AOC) curve, indicating behavioral criterion on individual sessions and their average (triangles) for test stimuli inside and opposite side of the RF during four attention conditions (#sessions: 20 monkey S; 22 monkey P). Lines connect four attention conditions within a session. Error bars, 95% confidence intervals).

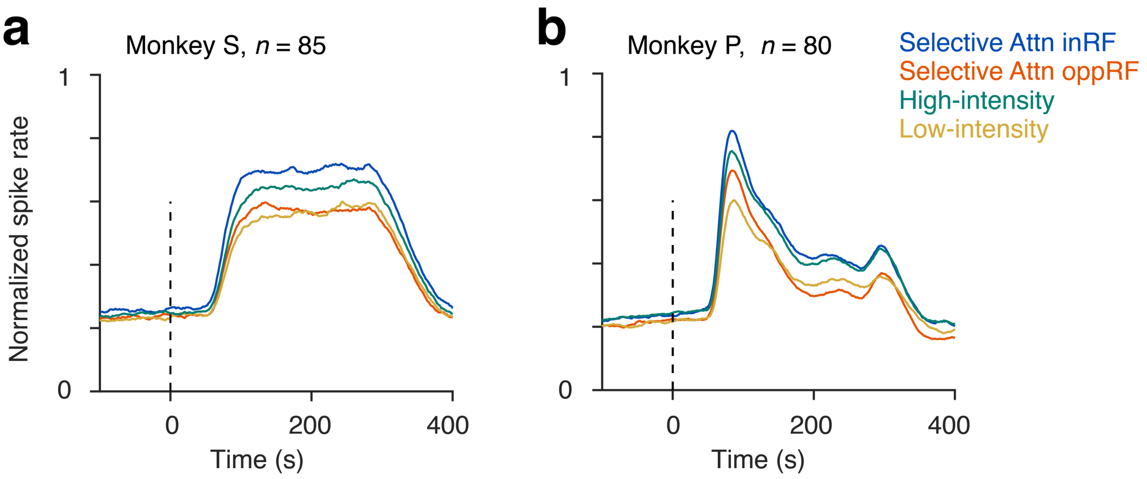


**Supplementary Figure S2. PSTHs of unique units across recording sessions.** A unique unit was randomly selected from every electrode out of a multielectrode array (96 channel) across session. Distribution of neuronal modulation (*d'*) of these unique units (n = 159) are plotted. Red bars, neurons with MI values significantly different from zero (nMI >0 = 99/ 159, nMI <0 = 3/ 159, p < 0.05). White bars, non-significant MI. Solid triangle, mean population MI across all unique units (n = 159, p < 10–25; t-test).

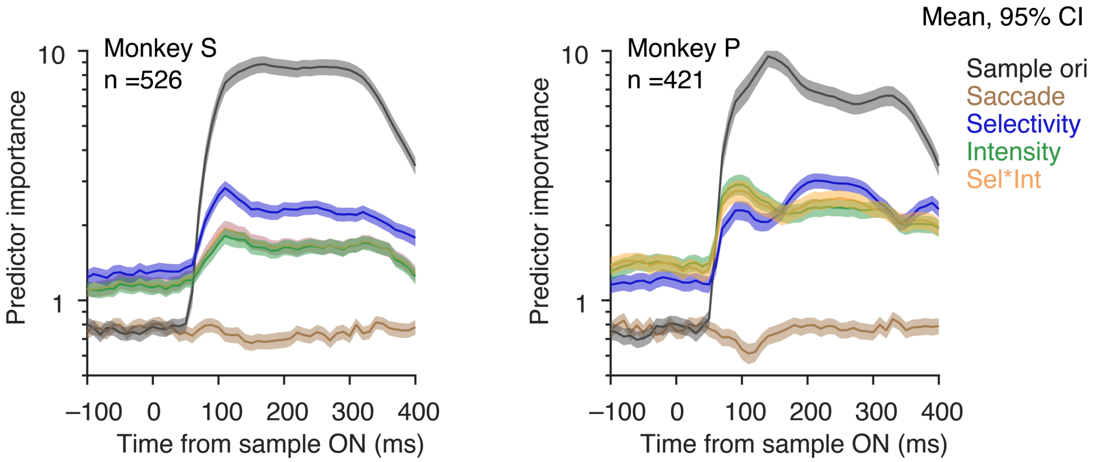


**Supplementary Figure S3. Encoding of attention components within single-trial spike trains: different task-demand context.** Comparing predictor importance (PI) that measures contributions of different predictor variables estimated by absolute standardized predictor coefficient values of recorded units from monkey S (left, n = 526) and monkey P (right, n = 421) that were significant fit with the generalized linear model in Figure 4. Error bars, 95% confidence intervals (bootstrap, $n = 10^4$).

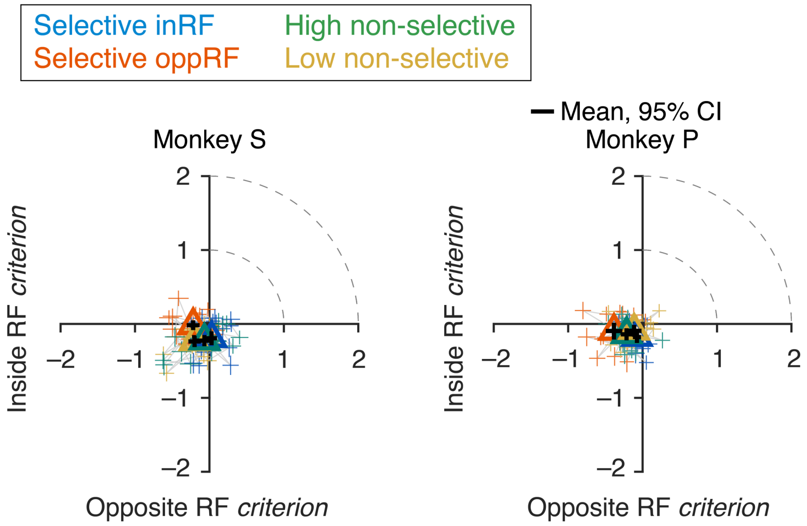


**Supplementary Figure S4. Behavioral criterions associated with attentional selectivity and intensity controlled by reward size.** Attention operating characteristic (AOC) curve, indicating behavioral criterion on individual sessions and their average (triangles) for test stimuli inside and opposite side of the RF during four attention conditions (#sessions: 20 monkey S; 16 monkey P). Lines connect four attention conditions within a session. Error bars, 95% confidence intervals).

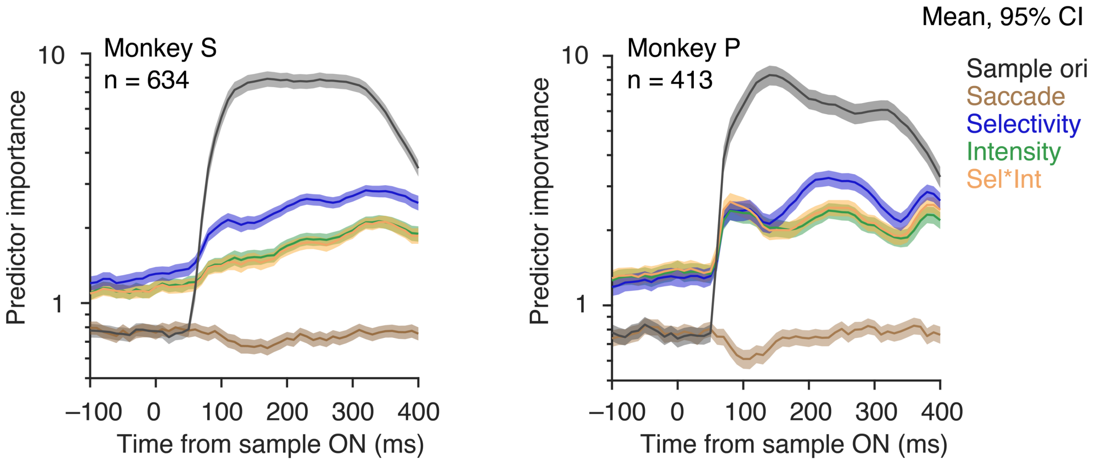


**Supplementary Figure S5. Encoding of attention components within single-trial spike trains: different reward context.** Comparing predictor importance (PI) that measures contributions of different predictor variables estimated by absolute standardized predictor coefficient values of recorded units from monkey S (left, n = 634) and monkey P (right, n = 413) that were significant fit with the generalized linear model in **Figure 7d-f**. Error bars, 95% confidence intervals (bootstrap, n = $10^4$).

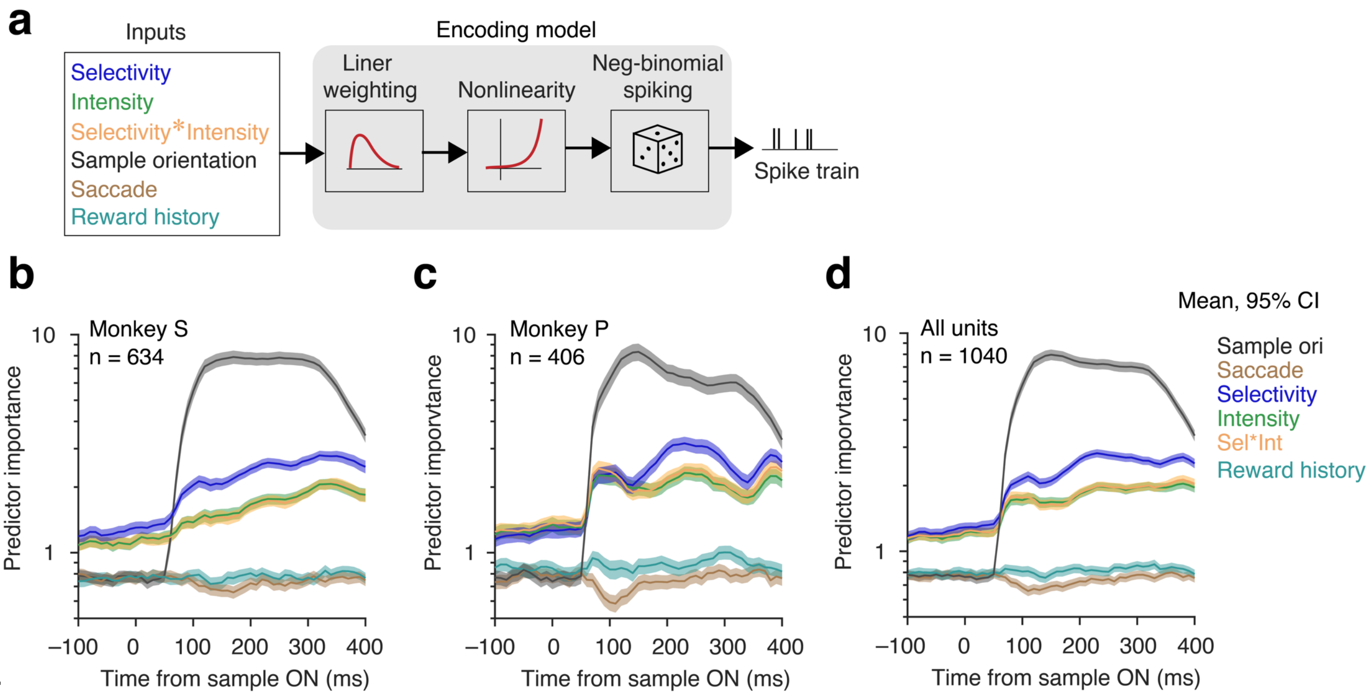


**Supplementary Figure S6. Alternate encoding model containing reward history: different reward context. b-d** Comparing predictor importance (PI) that measures contributions of different predictor variables estimated by absolute standardized predictor coefficient values of recorded units from monkey S (left, n = 634) and monkey P (right, n = 413) that were significant fit with the generalized linear model in **Figure 7d-f**. Error bars, 95% confidence intervals (bootstrap, n = $10^4$).

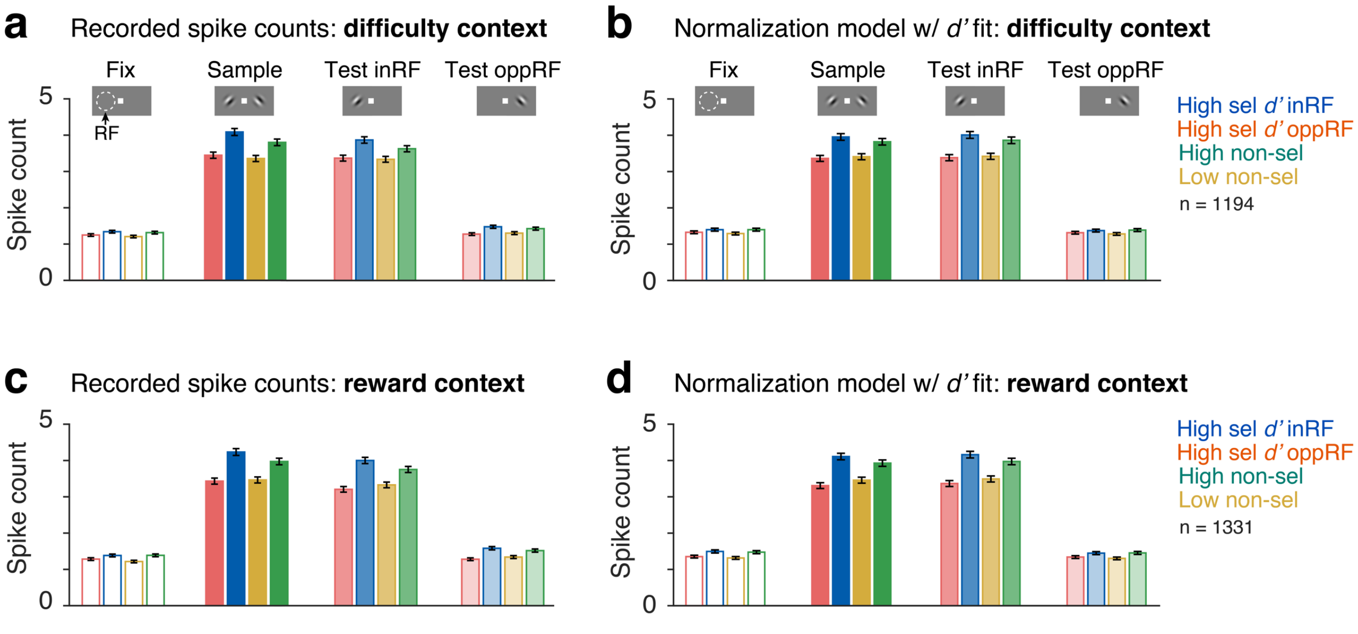


**Supplementary Figure S7. Comparing population averaged observed and normalization model fitted spike counts across different stimulus and attention conditions. a** Observed population averaged spike counts over 200 ms during pre-sample (–200 to 0 ms), sample stimuli (60 – 260 ms) and test stimulus (60 – 260 ms) periods for four selective and non-selective attention conditions in task difficulty sessions (n = 1194). **b** Same as in (a) for normalization model (model w/ *d'*) fitted spike counts. **c, d** Same as in (a) and (b), observed (c) and model fitted (d) spike counts in varying reward sessions (n = 1331).
